# The embryonic node functions as an instructive stem cell niche

**DOI:** 10.1101/2020.11.10.376913

**Authors:** Tatiana Solovieva, Hui-Chun Lu, Adam Moverley, Nicolas Plachta, Claudio D. Stern

## Abstract

In warm-blooded vertebrate embryos (mammals and birds), the body forms from a growth zone at the tail end. Hensen’s node, a region which induces and patterns the neural axis is located within this growth zone. The node also contains the precursors of neural, mesodermal and endodermal structures along the midline and has been suggested to contain a small population of resident stem cells. However, it is unknown whether the rest of the node constitutes an instructive stem cell niche, specifying stem cell behaviour. Here we combine transplantation of a single cell in vivo with single-cell mRNA sequencing in the chick and show that when made to enter the node, non-node-progenitor cells become resident and gain stem cell behaviour. These cells preferentially express G2/M phase cell-cycle related genes and are concentrated in posterior sub-regions of the node. The posterior part of the node therefore behaves as an instructive stem cell niche. These results demonstrate a new function for the vertebrate node during development.

---

In higher vertebrate embryos, the body axis forms in head-to-tail direction from a growth zone at the tail end which is present from gastrula stages through to the end of axis elongation. During gastrulation (in chick: stage HH3+ to HH4)^1^, epiblast cells lateral to the anterior tip of the streak/node ingress into it. After this, the node begins to regress caudally^2^ as cells exit the node to lay down the midline of the developing head-tail axis (Fig. 1 a-c)^3–6^. However, transplantation of cell groups and fate mapping experiments in chick^4,7–9^ and mouse^10–14^ during early development have suggested that the node contains some resident, self-renewing cells that persist during axial elongation in the node, while other cells leave (Fig. 1c, ‘RC’). Could the former be stem cells^15^, specified by neighbouring node cells?

**Figure 1.**
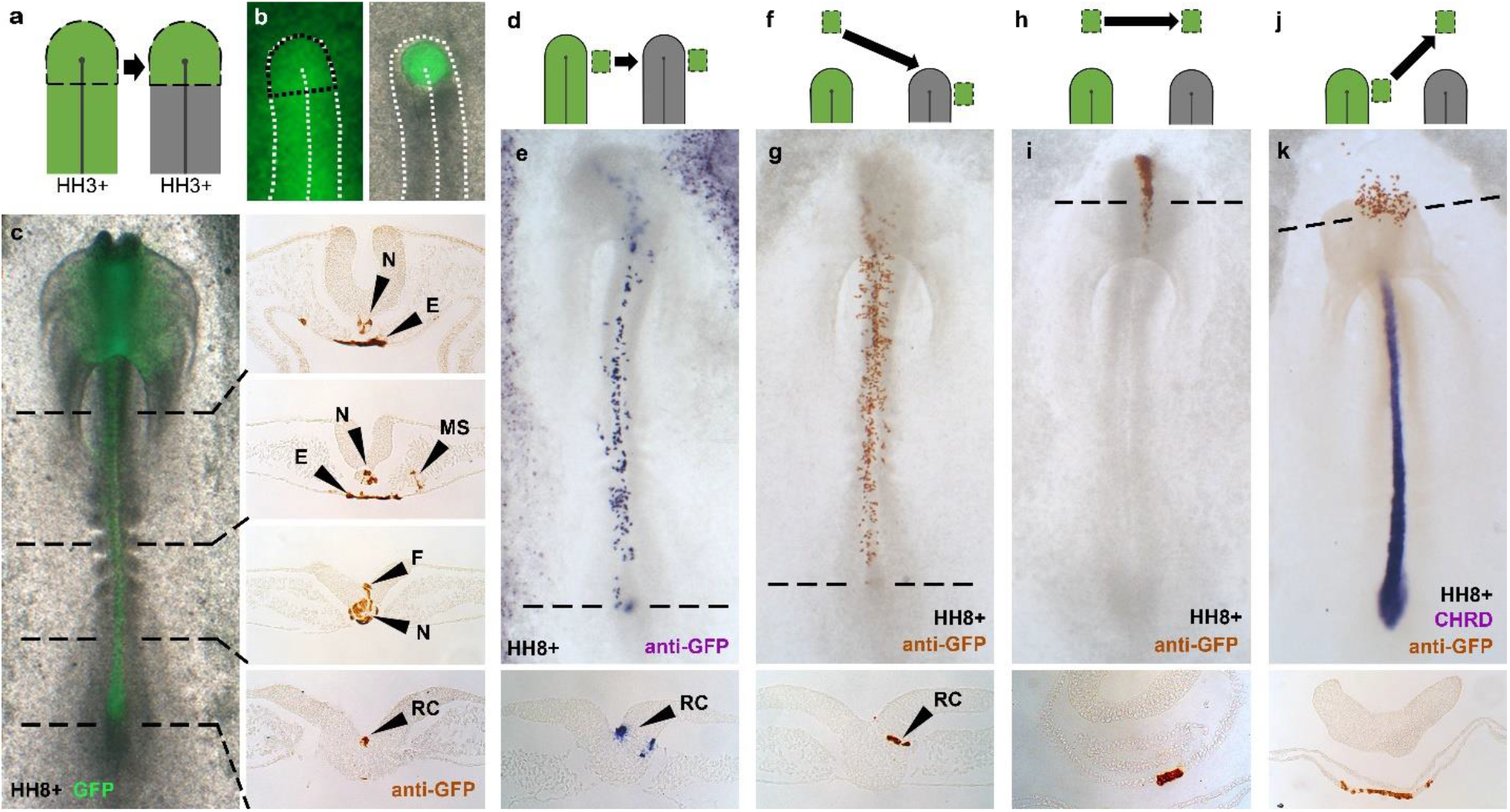
The node confers resident behaviour and axial fates. **a-c,** Node replacement using a GFP-donor showing normal node axial fates. **d-e,** Epiblast lateral to the HH3+/4 node ingresses into it and gives rise to the axis and to regressing node as ‘resident cells’ (RC). **f-g,** Anterior epiblast not normally fated to enter the node behaves as lateral epiblast when forced to do so. **h-i,** Anterior epiblast normally gives rise to head structures. **j-k,** Lateral epiblast no longer gives rise to node-derived axial structures when prevented from entering the node. N=notochord; E=endoderm; MS=medial-somite; F=floorplate; RC=resident cell. Transverse dashed lines show levels of accompanying sections.

There are two possibilities: either there is a special population of stem cells set aside during early development that is maintained by the node environment (“permissive”), or the node constitutes a special niche that can instruct any cell to acquire self-renewing stem cell characteristics^16–18^. To demonstrate self-renewal and to test whether the node is an instructive stem cell niche, it is critical to challenge the responses of an individual cell to the node environment. Here we use transplantation of single cells in vivo and single-cell RNA sequencing (scRNA-seq) to describe a cell’s response to the node.

## Non-node cells can become resident

To test whether the node environment can impart resident behaviour onto other cells, we grafted anterior epiblast (which never normally enters the node^5,19–21^) to a position adjacent to the HH3+/4 node, so that transplanted cells would be carried into the node by gastrulation movements (Fig. 1f). Graft-derived cells (from the transgenic GFP-donor) give rise to axial tissues and express appropriate molecular markers of node, notochord and somite (Fig. 1g, Extended Data Fig. 1a-j). Importantly, the contribution of this anterior epiblast to cells with resident behaviour (88%, n=30/34) is similar to that of lateral epiblast (Fig 1. d-e, Extended Data Fig. 1k) (89%, n=8/9), which does normally enter the node. These results show that the node can confer resident behaviour and axial identity.

## Prospective node cells are plastic

To test whether cells are intrinsically committed to node and axial identities, we prevented lateral epiblast cells from entering the node by grafting them into a remote anterior position (Fig. 1j). After culture to HH8-10, graft-derived cells localize to and resemble head structures rather than node derived tissues and fail to express the node marker, chordin (Fig. 1k). Lateral cells therefore develop according to their new anterior position^5,19–21^ (Fig. 1h-i), demonstrating that cells normally destined to give rise to node and axial identities are not committed to these before they enter the node.

## Self-renewal specified by a node niche

We then asked whether resident cells specified by the node are stem cells by testing for self-renewal, a key characteristic of a stem cell^22–24^. First, anterior epiblast was made to enter the node by grafting adjacent to it at HH3+/4 (Extended Data Fig. 2a-b). Following culture to HH8-10, two to ten GFP-positive cells remaining in the node were re-grafted into a second, younger (HH3+/4) host node (Extended Data Fig. 2c-f) to determine whether the GFP-positive cells can self-renew and contribute daughters to the developing axis for a second time. GFP-positive cells contributed to both node and axis in 17% of embryos (n=4/23) (Extended Data Fig. 2g-i) suggesting self-renewal.

To demonstrate this at the single-cell level we repeated the re-grafts (again using two successive hosts) with just a single GFP-positive resident cell (Fig. 2). After culture of the second host to HH8-10 (Fig. 2e-f), GFP-positive cells were detected in 21% of grafted embryos (n=11/53) (Extended Data Fig. 2j), of which 36% had multiple GFP-positive cells, showing that cell division had occurred; 18% had GFP-positive cells in both node and axis revealing that the resident cells both self-renewed and contributed to the axial midline (Fig. 2f-g). The node’s ability to specify self-renewing resident behaviour from cells not normally destined to enter it clearly demonstrates the properties of an instructive niche^16–18^.

**Figure 2.**
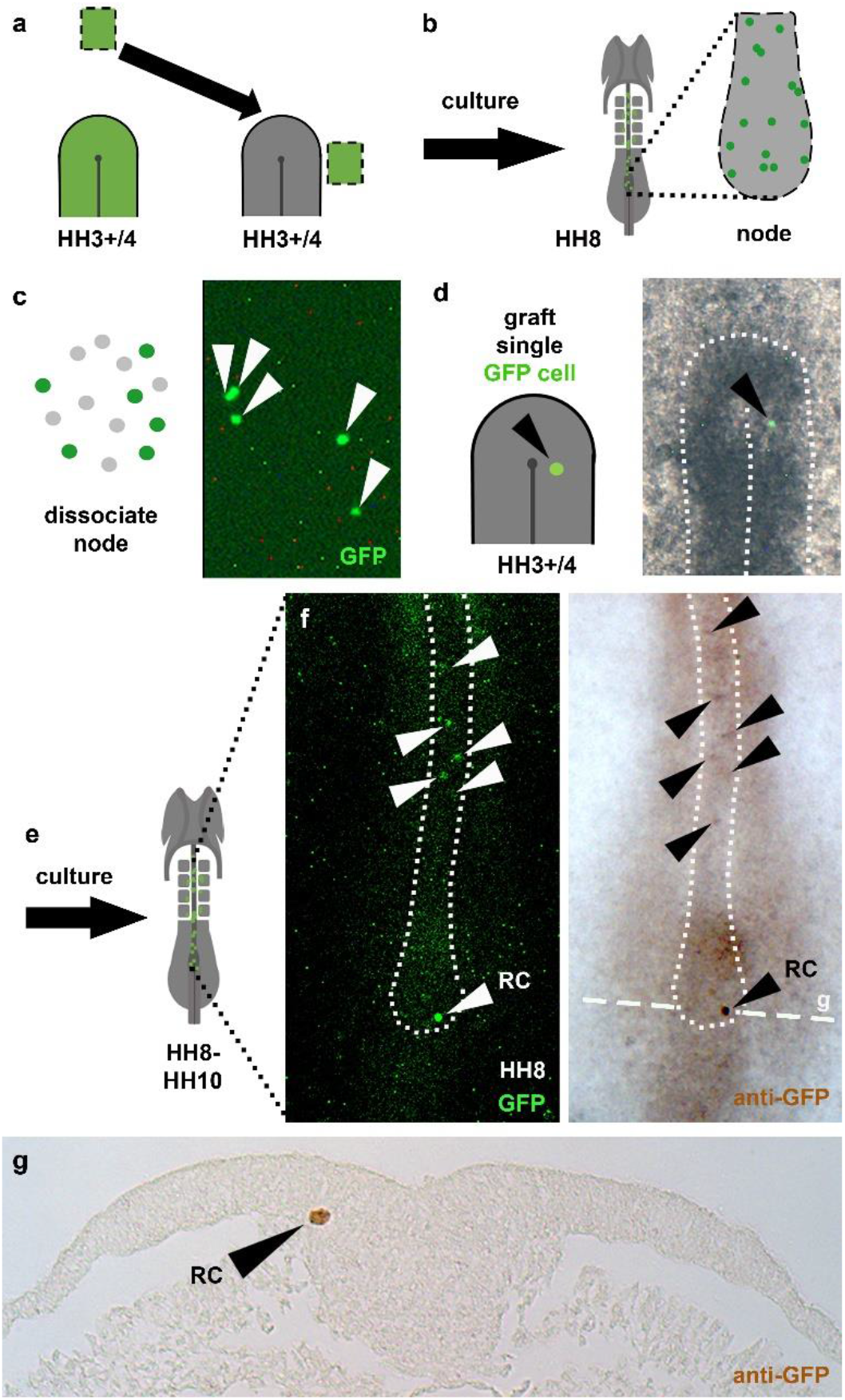
Single cell re-grafts reveal that the node can specify self-renewing resident cells. **a,** Anterior epiblast not normally fated to enter the node was made to do so. **a-c,** After culture, cells still residing in the HH8 node (b) were dissociated (c). **d,** A single GFP-positive cell was isolated and grafted into a second younger, HH3+/4 host node. **e-g,** After culture (e), the single GFP-positive cell had proliferated and descendants were found resident in the node and along the axis (f-g). Transverse dashed line in (f) shows level of section in (g). RC=resident cell.

To test whether this niche relies on neighbouring supporting cells, re-grafts of an individual GFP-positive cell were performed alone (n=26), or attached to a few neighbouring GFP-negative cells from the first host (n=27). While survival and division of GFP-positive cells was comparable between these two conditions, GFP-positive cells attached to GFP-negative neighbours showed an increased contribution to the node when compared to GFP-positive cells grafted alone (40% versus 17%) (Extended Data Fig. 2j). This finding is also consistent with the idea that the node behaves like an instructive niche, inducing self-renewal behaviour on other cells.

## Properties of node sub-regions

Does the entire node act as a niche, or is this property located in a particular sub-region? To identify the regions containing long-term resident cells, we constructed a fate map of the node by labelling each of six sub-regions using a lipophilic dye (DiI) at HH8 (Extended Data Fig. 3a). After culture to HH11-12, all cells from anterior sub-regions had come out from the node and the middle sub-regions contributed to anterior parts of the later node (chordoneural hinge), while only the posterior sub-regions continued to contribute to the entire older node and its derivatives (Fig. 3a-b, Extended Data Fig. 3b-g). This suggests that resident cells remaining in the node the longest are confined to posterior sub-regions. Consistent with this, in the regrafting experiments described earlier, the single cell remaining in the node was always found in the posterior sub-region (Fig. 2f-g). Furthermore, live imaging of embryos in which a mosaic of cells was fluorescently-labelled revealed endogenous resident cells remaining in the posterior node as it regresses (HH5-9), whereas most cells from other regions of the node were left behind to contribute to the axis (Supplementary Movie 1). This is consistent with fate mapping results of the mouse node, showing that some labelled cells remain in the node-primitive-streak border as the axis forms^14,25^. The posterior node is therefore the strongest candidate for an axial stem cell niche in amniotes. This is further supported by findings that when the posterior part of the node is removed, axial elongation is impaired^9,25^.

**Figure 3.**
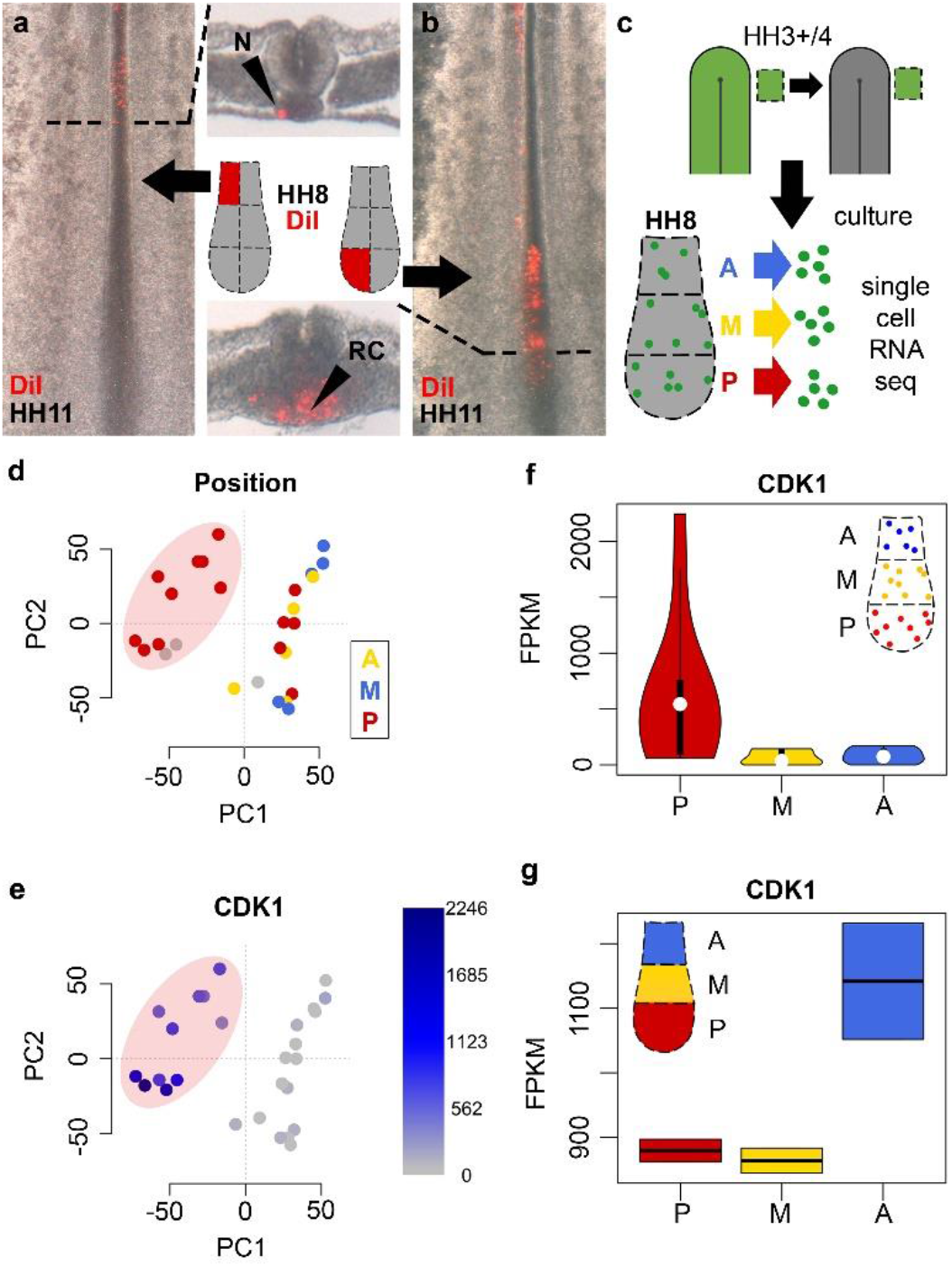
Long-term resident cells reside in the posterior node and are enriched in G2/M-phase cell-cycle genes. **a-b,** Fate mapping of six domains reveals that only posterior sub-regions continue to contribute to resident cells. **c,** Individual resident cells originating from lateral epiblast were isolated from anterior (blue), middle (yellow) and posterior (red) HH8 node sub-regions and processed for scRNA-seq. **d-e,** PC analysis reveals a cluster of posterior cells (pink oval) (d) enriched in G2/M-phase-related genes, including CDK1 (e) (intensity of blue reflects FPKM levels). **f-g,** While single posterior resident cells are enriched in CDK1 (f), bulk-RNA-seq shows that this is not the case in all cells of that region.

## Molecular properties of resident cells

What are the molecular characteristics of cells residing in this posterior niche? We grafted GFP-epiblast from next to the node to the same position in a wild-type host and cultured the embryos to HH8. Single graft-derived cells were collected from the posterior, middle and anterior regions of the HH8 node, and processed for scRNA-seq using SmartSeq (Fig. 3c). The data were examined by Principal Component (PC) analysis (Extended Data Fig. 5a); the first component (PC1) groups cells into two clusters, one composed largely of posterior cells (Fig. 3d). To identify the genes causing this clustering, we calculated the correlation coefficient of genes for PC1. Thirty-seven genes have significant expression in the posterior cluster (correlation coefficient <-0.8); the majority of these (31/37) encode proteins of the G2/M phases of the cell cycle (Fig. 3e-f, Extended Data Fig. 4). This suggests that these cells are preparing to divide. Graft-derived cells isolated from other parts of the node appear to be randomly distributed in other phases of the cell cycle (Extended Data Fig. 5b-f).

To determine whether the posterior region of the node is a unique site for cells at the G2/M phases of the cycle, we compared the transcriptomes of entire node sub-regions at HH8 (Extended Data Fig. 7a). Half of the cell-cycle related genes identified in the scRNA-seq were also enriched in the posterior node as a whole, while the other half, including CDK1, were instead enriched in anterior regions (Fig. 3f versus 3g, Extended Data Fig. 6). Expression of these G2/M cell-cycle related genes is therefore specific to individual resident cells, but not necessarily to their micro-environment. This suggests that dividing resident cells may make up only a small proportion of the node niche. Interestingly, the transcriptomes of the sub-regions reveal an enrichment of genes involved in Wnt, Notch and FGF signalling in the posterior part of the node (Extended Data Fig. 7). These three pathways have been implicated in other stem cell niches^18^.

## Competence to respond to the node niche

If anterior epiblast cells can acquire resident, self-renewing behaviour in response to the node environment, can any epiblast cell be similarly induced? To answer this, epiblast from older stage embryos (HH4+/5, corresponding to prospective neural plate) was used as donor tissue (Fig. 4a-b)^3,26^. This later HH4+/5 epiblast was forced to enter a younger (HH3+/4) node by grafting just adjacent to it (Fig. 4c-e). After culture to HH8-11, graft-derived cells from these late-to-early transplants contributed to the same axial and paraxial structures as the grafts from younger donors described earlier (Extended Data Fig. 8a). However, late epiblast gave rise to caudal node and/or chordoneural hinge in fewer embryos (55%) than lateral or anterior epiblast (89% and 88% respectively) suggesting that late epiblast cells are less able to respond to the node environment. Late epiblast cells also give rise to mesodermal structures (notochord and PSM/medial somite) less frequently than either lateral or anterior epiblast, and some that do, fail to express appropriate mesodermal genes (Fig. 4e). In contrast, late grafts contribute more frequently to neural structures (floorplate, 45% and lateral neural plate, 27%) than younger lateral epiblast (33% and 11% respectively) (Extended Data Fig. 8a). At the time of grafting, donor late epiblast cells already express neural plate markers (including SOX2 and ZEB2) (Extended Data Fig. 8b-g), but following grafting they lose expression of these genes except for descendants contributing to neural structures (Extended Data Fig. 8h-k). This suggests that the node environment causes late epiblast cells to lose their neural plate identity but is not sufficient to convert them fully into axial mesoderm. This transition away from mesodermal and towards neural fates appears to take place around stages HH5-to HH5 (Extended Data Fig. 8l) and suggests that older, neural plate epiblast has lost its competence to respond to signals from the node niche that induce axial identity and resident behaviour.

**Figure 4:**
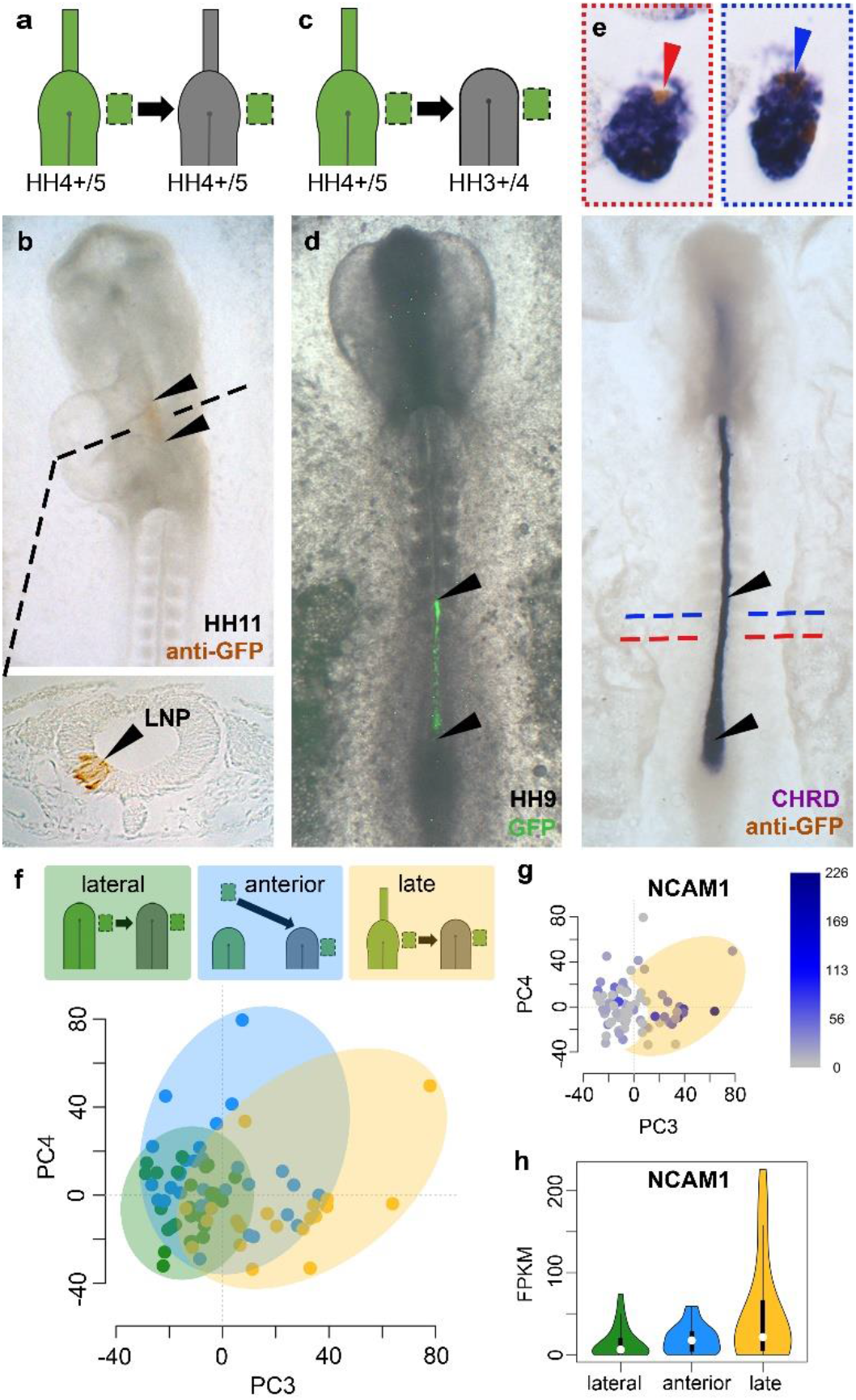
Older epiblast is not competent to respond to the node. **a-b,** (‘Late’) epiblast lateral to the HH4+/5 node normally contributes to lateral neural plate (LNP). **c-e,** When made to enter the younger node (c), ‘ late’ epiblast contributes to the axial midline (d-e). Black arrowheads show the extent of head-to-tail contribution. While some graft-derived cells (visualised by anti-GFP) in the notochord express the notochord marker CHRD (blue arrow), others do not (red arrow). **f-h,** Single cells from the HH8 node plotted using PC3/4 cluster according to their epiblast origin (lateral, anterior or late) (f). Expression of neural plate-like genes, including NCAM1, correlates with ‘late’ cells outside of the control (green) cluster (yellow oval) (g-h). Intensity of blue in (g) reflects FPKM levels.

What molecular changes underlie this loss of competence? To address this we performed scRNA-seq on cells with resident behaviour at HH8, originating from non-competent cells (from late-to-early grafts) and compared these to cells that are competent (from grafts of lateral or anterior epiblast) (Extended Data Fig. 9a-c). Irrespective of origin, the most significant variation among the cells could still be accounted for by their expression of G2/M-phase related cell-cycle genes (Extended Data Fig. 9d-g, compare with Fig. 3d-e; clustering by PC1-2). However, PC3-4 clusters cells into overlapping groups according to their donor origin (lateral, anterior or late epiblast) (Fig. 4f). A subset of late-epiblast-derived cells that is associated with limited competence is characterised by specific expression of 15 genes (correlation coefficient >0.55 with PC3) (Extended Data Fig. 10a). Of these genes, seven have known roles in cell adhesion and/or in neural development including NCAM1 and CLDN1, normally expressed mainly in the neural plate (Fig. 4g-i and Extended Data Fig. 10b-i). This suggests that these later cells may have lost their competence to respond to the node environment because they have already initiated their differentiation into neural plate.

## Conclusion

Until now, apart from a few studies using prospective single cell fate mapping^4,10,11^, evidence for resident cells in the node was based on studies of cell populations^8,9,12–14^. Without challenging behaviour at the single cell level it has not been possible to test whether the node represents an instructive stem cell niche. Here, using single cell grafts we discovered that the node can specify resident and self-renewing behaviour. Thus, in addition to its well-known roles as an ‘organizer’ of the amniote embryo^27–29^ and its ability to dorsalize mesoderm^30,31^, this key embryonic structure also functions as a stem cell niche that can specify resident stem cells for the developing head-tail axis.

## Supporting information

RNAseq dataset

single cell RNAseq dataset

## Methods

### Embryos

Wild type chicken embryos were obtained from Brown Bovan Gold hens (Henry Stewart Farm). Transgenic cytoplasmic GFP chicken embryos were supplied by the avian transgenic facility at The Roslin Institute, Edinburgh^8^. All eggs were incubated at 38°C in humidified incubators and staged according to Hamburger and Hamilton^1^. For grafting experiments, *ex-ovo* embryo cultures were prepared using the New technique^32^ with modifications as described by Stern and Ireland^33^.

### Epiblast grafts

Donor embryos were isolated in Tyrode’s solution^34^. The donor embryo was turned ventral side up, underlying endodermal and mesodermal layers were peeled away and a piece of epiblast (~20-50 cells) was cut out using 30G syringe needles. The epiblast piece was checked to ensure no mesodermal/endodermal cells remained. An equal sized piece of epiblast was removed from the host in the desired location and replaced with the donor epiblast. For ‘lateral-to-lateral’ grafts (see Fig. 1d), epiblast was grafted into the equivalent position in the host as its donor origin (i.e. ‘left-to-left’ or ‘right-to-right’). The ‘lateral epiblast’ was taken from immediately adjacent to the tip of the streak/node. ‘Anterior’ epiblast was taken from a midline position, about half-way between the tip of the streak/node and the anterior area opaca.

### Re-grafts of groups of cells

The first graft was an epiblast graft from a GFP-donor to a non-GFP host (see Extended Data Fig. 2a-b). The second graft (re-graft) included a group of cells from the first host’s node, containing 2-10 GFP-positive cells alongside some neighbouring GFP-negative cells. A small ‘nick’ was made in the node (ventral side) of the second host into which this group of cells was inserted using 30G syringe needles to carefully manoeuvre the small pieces of tissue (see Extended Data Fig. 2c-f). The grafted embryos were left at room temperature for ~15 min to aid attachment of the graft to the host before further incubation. Embryos were cultured to HH8-10.

### Single cell re-grafts

The first host had an ‘anterior-to-lateral’ graft (see Fig. 2a). After culture to HH8-10, a single GFP-positive cell was collected from the host node (see ‘single cell manipulation’ below) and then transferred using a micropipette made from a pulled 50 μl calibrated micropipette (Drummond Scientific, Cat 2-000-050) attached to an aspirator tube, into the second host (HH3+/4) (see Fig. 2b-d). For some re-grafts, a single GFP-positive cell was transferred attached to one or more neighbouring GFP-negative cells from the first host (but there was never more than one GFP-positive cell). A small ‘nick’ was made in the node of the second host. The GFP-positive cell was maneuvered into this nick by gently ‘blowing’ saline on the cell with a micropipette. Ideally, once placed into its pocket, a flap of tissue would be used to cover the transplant site. The grafted embryo was then left at room temperature for ~15 min to aid attachment of the cell to the host. Each New culture was checked by fluorescence microscopy again just prior to incubation to ensure that the grafted GFP-positive cell was still in place. Embryos were cultured to HH8-10.

### DiI labelling

The lipophilic dye, DiI (DiI-CellTracker CM, Molecular Probes Life Technologies, # C7001) was used for fate mapping of the HH8 node. For 10 μl of working solution, 8.5 μl of 0.3M sucrose and 1 μl of 1:20,000 Tween-20 were used with 0.5 μl of 2mM DiI (in dimethylformamide). All components were first pre-heated at 65°C, thoroughly mixed and dissolved. The protocol for preparation and application of DiI was adapted from^4,35,36^. The embryo was first prepared for New culture^32^ and kept submerged in Tyrode’s. The node subregion to be labelled was cut out using 30G syringe needles and transferred to a drop of Tyrode’s containing DiI (~9:1 Tyrode’s: DiI working solution) and kept in the dark for 1-2 min. The tissue piece was then removed and washed in successive drops of Tyrode’s to remove any excess DiI before verifying that sufficient labelling had taken place, by fluorescence microscopy. The tissue piece was replaced into its original position, preserving the original dorsoventral orientation. Labelled embryos were cultured to ~HH11-12. After culture, embryos were fixed in 4% paraformaldehyde in PBS for at least 4 days at 4°C. To assess the location of the descendants of the DiI-labelled cells, several thick transverse sections were cut from each embryo by hand, using a scalpel, with the embryo pinned securely using insect pins in a silicon rubber-bottomed dish^34^.

### *In situ* hybridization

*in situ* hybridization with digoxigenin (DIG-)-labelled riboprobes was carried out following established protocols^37–39^. Antisense DIG-riboprobes were synthesized by restriction digest and *in vitro* transcription. Plasmids used: CHRD^40^, FOXA2^41,42^; PARAXIS/TCF-15^43^; ZEB2^26^; SOX2^44^; TBX6^45^; DLL1^46^; RSPO3 (ChEST784h18); NKAIN4 (ChEST110n2); DRAXIN (ChEST545l1); CLDN1 (ChEST168n2); NCAM1 (ChEST845i20); MSGN1 (ChEST90p23); CHST15 (ChEST391h17); AKAP12 (ChEST376j15); FOXM1 (ChEST313o15); TOP2A (ChEST849a2); NUF2 (ChEST450j22); CENPL (ChEST97i12); MAD2L1BP (ChEST365n5).

### Immunohistochemistry

anti-GFP antibody staining largely followed the methods described by Stern^38^ and Streit and Stern^39^. Embryos were processed for anti-GFP antibody staining either immediately after collection (and fixing in 4% PFA overnight at 4°C) or following *in situ* hybridization.

### Histology

Some embryos processed for *in situ* hybridization and/or anti-GFP antibody staining were embedded in paraffin wax and sectioned using a microtome. Methods largely followed those of Izpisúa-Belmonte et al.^37^. All sections were transverse and 10 μM thick. Slides were mounted using a 3:1 solution of Canada balsam (Merck, # 1016910100) and Histoclear (HS-202 HISTO-CLEAR II, National Diagnostics).

### Microphotography

Images of all whole-mount embryos and thick sections were recorded using transmitted light with an Olympus SZH10 stereomicroscope with epifluorescence optics. Paraffin sections were examined on an Olympus Vanox-T optical microscope. A QImaging Retiga 2000R Fast 1394 camera and QCapture Pro software was used for all image capture.

### Live imaging and cell tracking

Electroporation mixture containing 1 mg.ml^-1^ pDsRed-Express plasmid, 6% (wt/vol) sucrose and 0.04% (wt/vol) Fast Green FCF was applied dorsally, just lateral to the node of HH4-embryos to transfect ingressing cells. Electroporation was performed in a custom-made chamber with four pulses of 5 V, 50 msec width, 500 ms interval. Embryos were then cultured using a modification of New’s method^32,33^ in 35 mm plastic dishes with a glass coverslip base, and imaged with a Zeiss LSM 880 inverted microscope using a Plan-Apochromat 20x, 0.8 NA objective. Images were acquired at 10 min intervals using 3×5 tiling (10% overlap) to achieve coverage of the whole embryo. Image analysis and cell tracking were performed using Imaris (Bitplane) software. The embryo was imaged from the epiblast (dorsal) side but the output from Imaris is displayed as a mirror image (pseudo-ventral view).

### Single cell manipulation

For collection of single cells for single cell re-grafts and scRNA-seq, the cultured embryo was first submerged in Tyrode’s solution (for re-grafts) or sterile molecular grade PBS with 0.1% glucose (for scRNA-seq). The node was then divided into anterior, middle and posterior sub-regions of equal rostro-caudal length. Each of these regions containing GFP-positive cells was cut out, in turn, and placed in a drop of non-enzymatic dissociation medium (Sigma, # C5914-100ML), kept over ice. Each piece was washed twice in drops (~30 μl) of this dissociation medium while over ice. To help with dissociation, after ~5 min, the tissue was gently aspirated up and down using a micropipette (made from a pulled 50 μl calibrated micropipette (Drummond Scientific, Cat 2-000-050) attached to an aspirator tube). The micropipette was broken at the tip to have a diameter just narrower than the width of the tissue piece. Once the piece of tissue was fragmented, a capillary with a narrower tip was used for further dissociation to single cells in suspension. GFP-positive cells were identified by fluorescence under a dissection microscope (x70 magnification) and were individually aspirated using a micropipette. The cell was transferred into a drop of Tyrode’s (for re-grafts) or of sterile molecular grade PBS (for scRNA-seq) to replace the dissociation medium and to verify that there was only a single GFP-positive cell. Once verified, the cell was transferred (using a fresh pulled micropipette) to the second host (for re-grafts) or into a 200 μl tube containing 5 μl of lysis buffer and 5% RNase inhibitor (for scRNA-seq) (lysis buffer and RNase inhibitor from SMART-Seq v4 Ultra Low Input RNA Kit, Takara, # 634892). Once the dissociation process began, cells were collected for ~20 min, after which time any remaining dissociated tissue was discarded, and a new tissue piece taken from the embryo.

### scRNA-seq

The SMART-Seq v4 Ultra Low Input RNA Kit (Takara, # 634892) targeting mRNA, was used for preparing the single cells for sequencing. Amplified cDNA was purified using AMPure magnetic purification beads (Agencourt AMPure XP, Beckman Coulter #A63880). The DNA concentration of purified cDNA was checked by Qubit (dsDNA HS Assay Kit, Thermofisher # Q33230). All samples yielding at least 5 ng of cDNA were sheared by sonication (Covaris, S220/E220 focused-ultrasonicator, settings set to: 10% duty factor, 200 cycles per burst, 120 second treatment time, 175W peak incident power) to obtain ~500bp fragments for library preparation. The ThruPLEX DNA-seq, Dual Index Kit (Takara, # R400406) was used to construct dual indexed libraries for each sample. Libraries were individually purified, using AMPure magnetic purification beads (Agencourt AMPure XP, Beckman Coulter # A63880). DNA concentration of purified libraries was checked by Qubit (dsDNA HS Assay Kit, Thermofisher # Q33230) and size distribution of cDNA measured using Tapestation (Agilent High sensitivity D1000 screen tape, # 5067-5584). All libraries were individually diluted to 10 nM in elution buffer before pooling together. Pooled libraries were sequenced by UCL Genomics using an Illumina NextSeq sequencer with a 75bp single end read cycle kit. The average number of reads per cell was ~10 Million (range: 6,838,400-15,404,851). The single cell RNA-seq raw data have been deposited in EBI Array Express (accession number E-MTAB-9116).

### RNA-seq of tissues

Tissues (HH8 node sub-regions) were isolated from transgenic-GFP embryos using 30G syringe needles in sterile molecular grade PBS. Tissues were collected into RNAlater (Invitrogen, # AM7020). For each sample, tissues were collected from thirteen to seventeen embryos. RNA was extracted using the Micro Total RNA Isolation Kit (Invitrogen, # AM1931) and concentration and quality measured using Tapestation (Agilent High sensitivity RNA screen tape, # 5067-5579). The NEBNext Single Cell/Low Input RNA Library Prep Kit for Illumina (# E6420) was used for cDNA and library synthesis (performed by UCL Genomics). Libraries were sequenced by Illumina NextSeq using a 75bp single end read cycle kit. The average number of reads per sample (node sub-region) was ~22 Million (range: 19,572,310 to 24,523,360). The bulk RNA-seq raw data have been deposited in EBI Array Express (accession number E-MTAB-9115).

### RNA-seq data processing

Raw data were checked using FastQC^47^ to assess overall quality. Cutadapt^48^ was used to remove low-quality bases (Phred quality score <20) at the 3’ and 5’ ends, adapter sequences, primer sequences, and poly-A tails of each read. Reads were aligned to the galGal6 chicken genome using TopHat2^49^, alignment rates were 91.9%±0.3% (for scRNA-seq) and 86.3%±0.65% (for RNA-seq of tissues). Transcripts were counted and normalized using Cufflinks^50^ programs *cuffquant* and *cuffnorm* respectively. Data analysis was performed in the R environment (R-3.5.1).

For scRNA-seq, all sequenced cells passed quality control. The matrix of transcript FPKMs (Fragments Per Kilobase of transcript per Million mapped reads) contains expression of 24,353 genes in 77 samples (cells). Of these, 13,817 are expressed (with an FPKM >0.5) in at least two cells in our data. The top 5000 most variable of these genes were used for principal component (PC) analysis. PC analysis was carried out on two datasets: data from single cells from the HH8 node originating only from lateral epiblast (n=27) and data from single cells from the HH8 node originating from anterior, lateral and late epiblast (n=77). Correlation coefficients were calculated for the relationship between gene expression and a given principal component.

For RNA-seq of tissues, all node sub-regions passed quality control. Mitochondrial RNAs, ribosomal RNAs and microRNAs were excluded. Fold changes for each gene in posterior sub-regions against its expression in all other regions were calculated to find genes with the most marked differential expression between sub-regions (see Extended Data Fig. 7b).

### Data availability

The scRNA-seq and bulk-RNA-seq raw datasets generated during the current study are available in the EBI Array Express (https://www.ebi.ac.uk/arrayexpress/). Accession numbers: E-MTAB-9116 and E-MTAB-9115.

## Acknowledgements

We are grateful to Nidia De Oliveira for technical help, Claire Anderson, Jun Ong, Hyung Chul Lee and Irene De Almeida for providing some probes, to Kathy Niakan, Claudia Gerri and Norah Fogarty for guidance on scRNA-seq and help with the Covaris shearing (done at the Francis Crick Institute, London) and to James Briscoe and Andrea Streit for helpful comments on the manuscript.

## Author contributions

H-CL processed raw RNA-seq data, AM and NP made and analysed the movie. TS performed all other experiments and analysis. CDS coordinated the research and obtained the funding. TS and CDS designed the project and wrote the manuscript.

## Supplementary information

is available for this paper.

## Materials and correspondence

Correspondence and requests for materials should be addressed to Claudio D. Stern.

## Supplementary Information

**Supplementary Movie**

**Supplementary movie 1 | Live tracking of cells in the node from HH5 to HH9.** Time-lapse movie showing a mosaic of cells labelled with DsRed (pseudo-colour encoded as green). Selected cells originating from the anterior part of the node at HH5 are highlighted in blue and some originating from the posterior part of the node are highlighted in red. The outline of the node is highlighted by a white dashed line.

**Supplementary Data**

**Supplementary data 1 | FPKM table for scRNA-seq data.** Transcriptional profiling of single cells individually harvested from the HH8 node, originating from the three experimental conditions described in Fig. 4f. Details on the epiblast origin (developmental stages and position in the donor) of the cells and their positions within the host node at the time of collection are included in the raw data submitted to EBI Array Express (accession number E-MTAB-9116).

**Supplementary data 2 | FPKM table for bulk-RNA-seq data.** Transcriptional profiling of 6 sub-regions of the HH8 node (see Extended Data Fig. 7a). The raw data were submitted to EBI Array Express (accession number E-MTAB-9115). Abbreviations for the node regions at stage 8: S15_AL8: anterior-left; S16_AR8: anterior-right; S17_ML8: middle-left; S18_MR8: middle-right; S19_PL8: posterior-left; S20_PR8: posterior-right.

