## Extended data figures for "The embryonic node functions as an instructive stem cell niche"

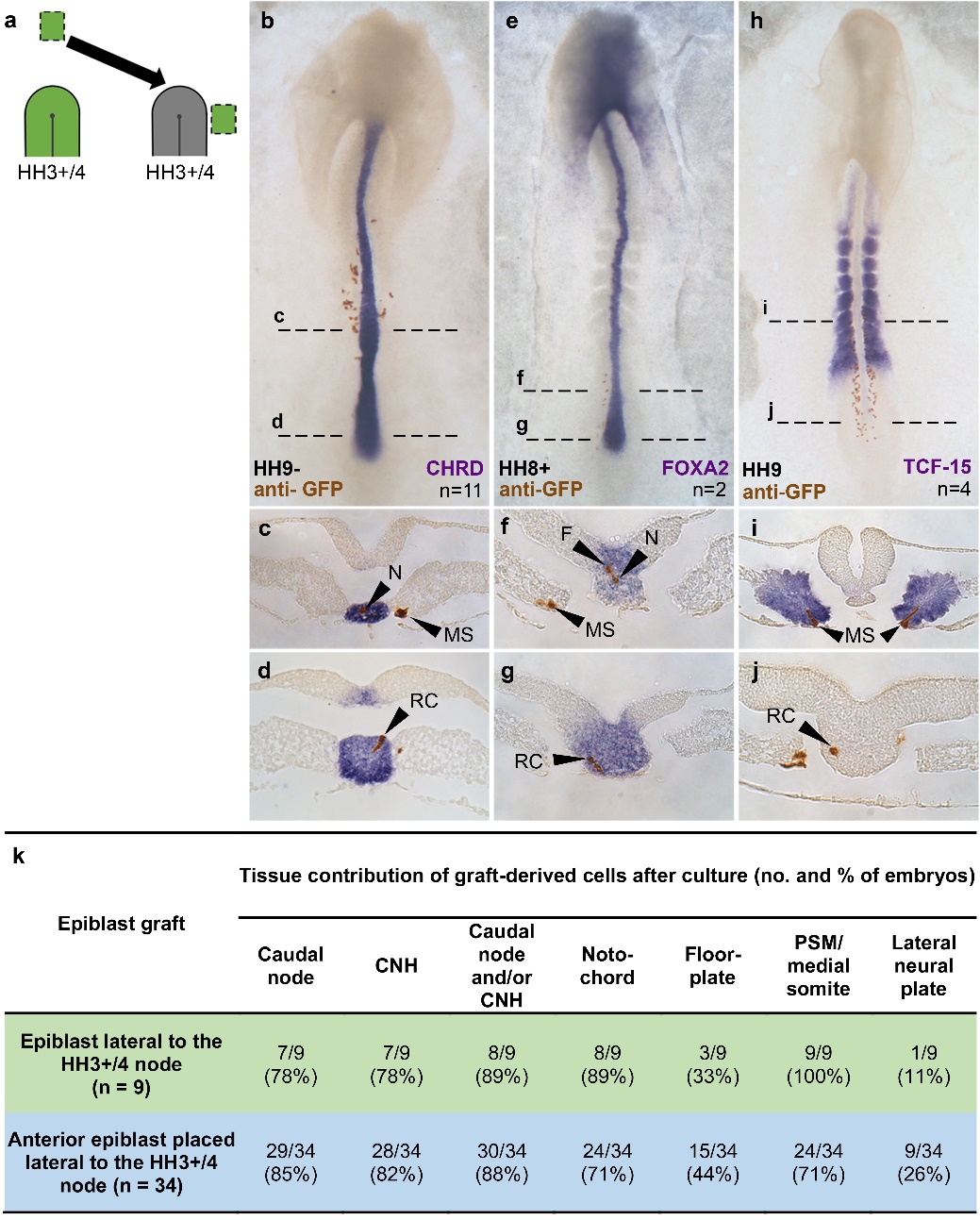

**Extended Data Figure 1 | Anterior epiblast derived cells made to enter the node express genes appropriate to their new location. a,** Schematic of graft using anterior epiblast from a GFP donor. **b-j,** Resulting embryos after culture to HH8-10, processed for *in situ* hybridization for the node and notochord markers CHRD (**b-d**) and FOXA2 (**e-g**) and the somite marker TCF-15/paraxis (**h-j**) and stained for anti-GFP antibody (brown). All whole-mount embryos shown in ventral view. **k**, Table of graft derived tissue contributions from lateral (green) and anterior (blue) epiblast grafts. n-numbers indicate the number of embryos grafted. N = notochord, MS = medial somite, F = floorplate, RC = resident cell, CNH = chordoneural hinge, PSM = presomitic mesoderm.

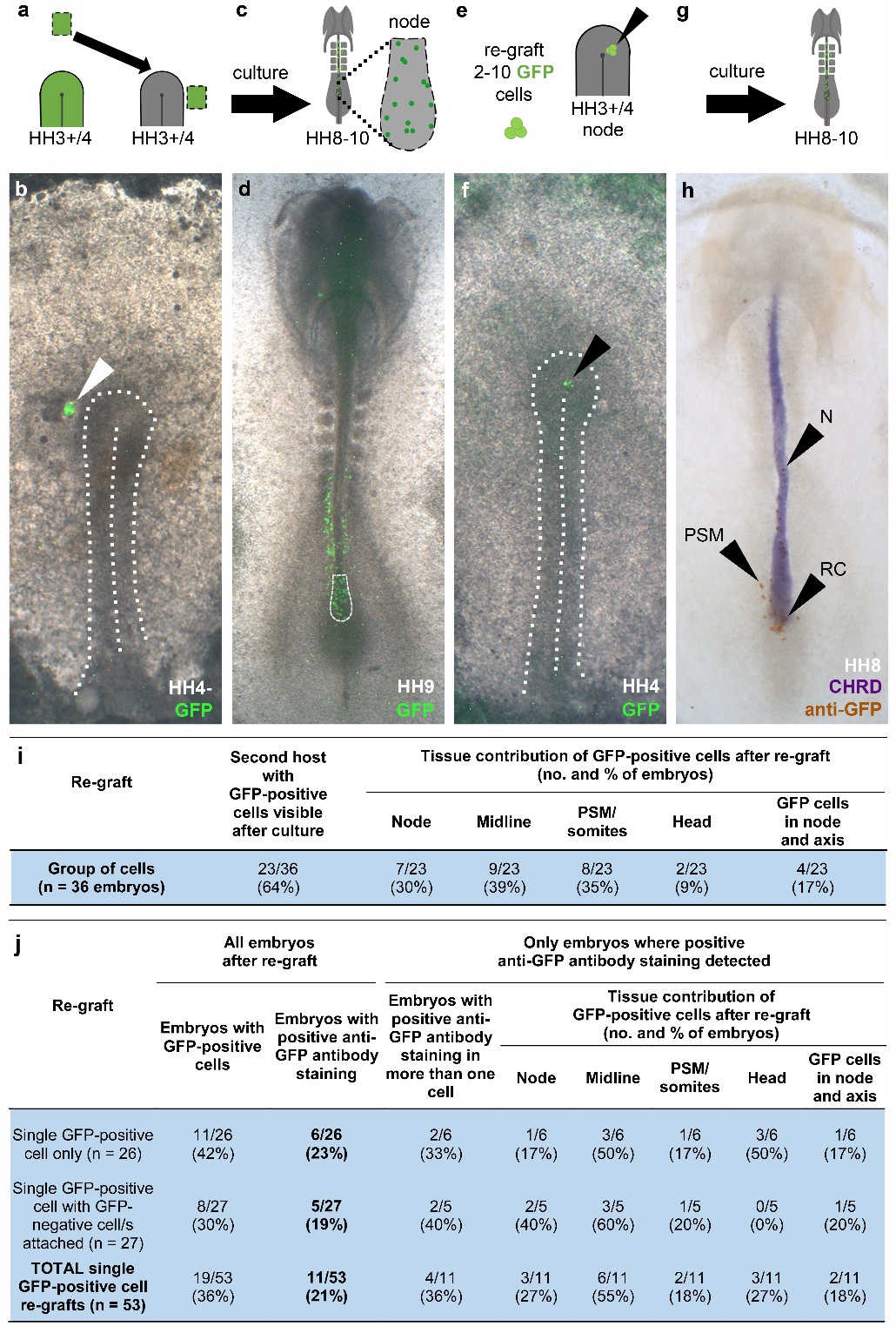

**Extended Data Figure 2 | Resident cells specified by the node can contribute to the node and axial midline for a second time. a-h**, Re-grafts of groups of GFP-positive cells with resident behaviour. First, anterior epiblast was grafted just lateral to the HH3+/4 node (**a-b**). (White arrow = GFP-cells in host). Embryos were then cultured to HH8-10 (**c-d**). Two-to-ten GFP-positive cells were taken from the regressing node (outlined in (**d**)), attached to some non-GFP neighbours and transferred to the node of a second, younger, HH3+/4 host (**e-f**). (Black arrows in E-F mark the re-grafted group of cells). After culture to HH8-10, GFP-positive cells can be found in both node and the midline (**g-h**). Black arrows in (**h**) point to some of the GFP-positive cells that contributed to notochord (N), presomitic mesoderm (PSM) and resident cells in the node (RC). (**b)** and (**d)** correspond to the same embryo; (**f)** and (**h)** are also two views of the same embryo. Cells re-grafted into the embryo in (**f)** were taken from the embryo shown in (**d)**. All embryos shown in ventral view. **i,** Summary table showing tissue contributions to the second host from re-grafted groups of GFP-positive resident cells (as in **h**). **j,** Summary table showing tissue contributions to second host from re-grafted single GFP-positive resident cells (as in Fig 2f). Single cells for re-grafting all came from primary grafts that used anterior donor epiblast (which would normally never enter the node), as shown in Fig. 2a. n-numbers refer to number of embryos grafted.

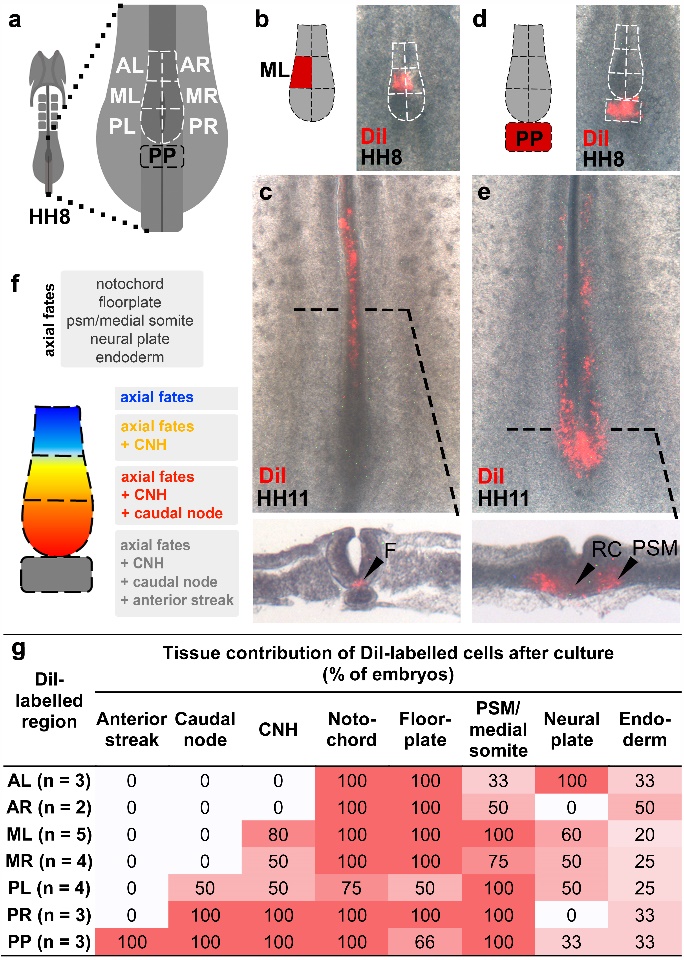

**Extended Data Figure 3 | DiI fate mapping of HH8 node/anterior streak sub-regions. H**H8 node outlined with white-dashed line and sub-divided into six sub-regions (ventral view) (**a**): anterior left (AL), anterior right (AR), middle left (ML), middle right (MR), posterior left (PL), posterior right (PR), anterior streak just caudal to the posterior part of the node (PP). Example of labelling a middle sub-region (**b**) and resulting embryo after culture (**c**). Example of labelling the anterior streak (**d**) and resulting embryo after culture (**e**). Summary of node sub-region fates illustrated in (**f**) and shown in a table in (**g**). n-numbers refer to the number of embryos labelled. The intensity of the red background reflects the percentage of embryos with labelled cells in the region shown in each column. F, floorplate; PSM, presomitic mesoderm; RC, resident cell.

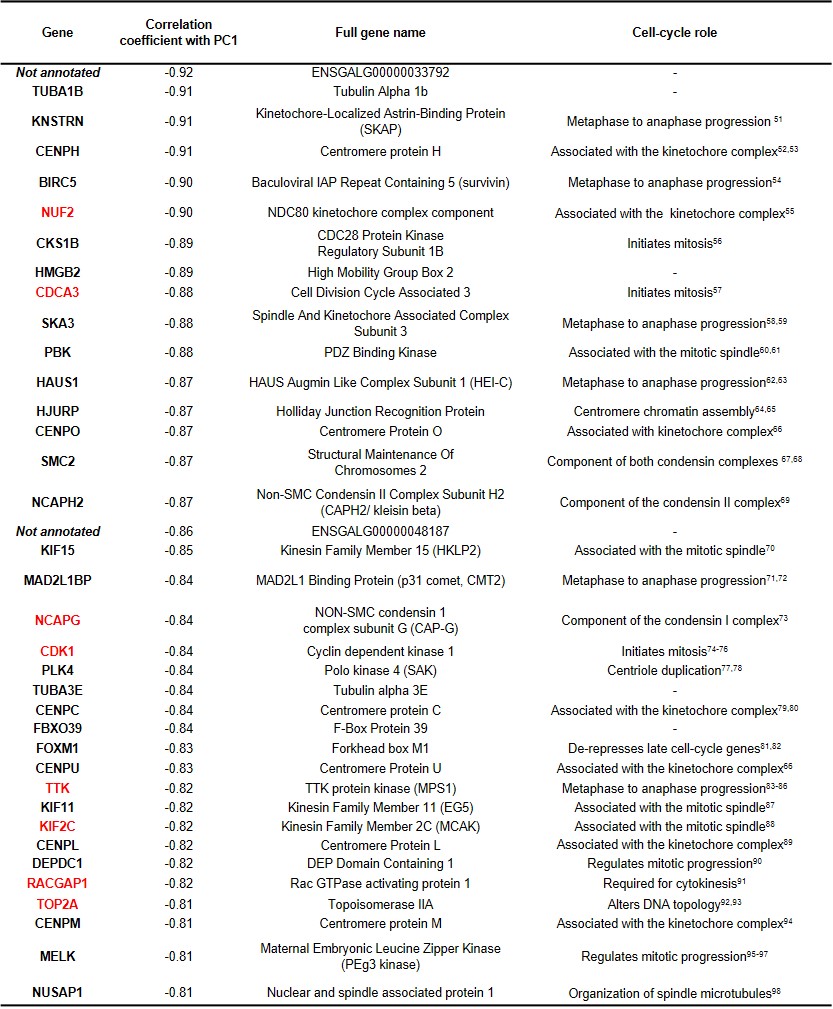
**Extended Data Figure 4 | Cells of the ‘posterior cluster’ preferentially express genes involved in G2/M phases of the cell cycle.** Correlation of gene expression with PC1 from scRNA-seq data. Of 37 genes with a correlation coefficient <0.80 for PC1, at least 31 are involved in G2/M phases of the cell cycle. Key cell-cycle-related roles of each gene outlined in column 4. Genes in red have been reported to be regulated by FOXM1, a specific transcriptional activator of G2/M phase related genes (which is itself represented among the genes in this group).

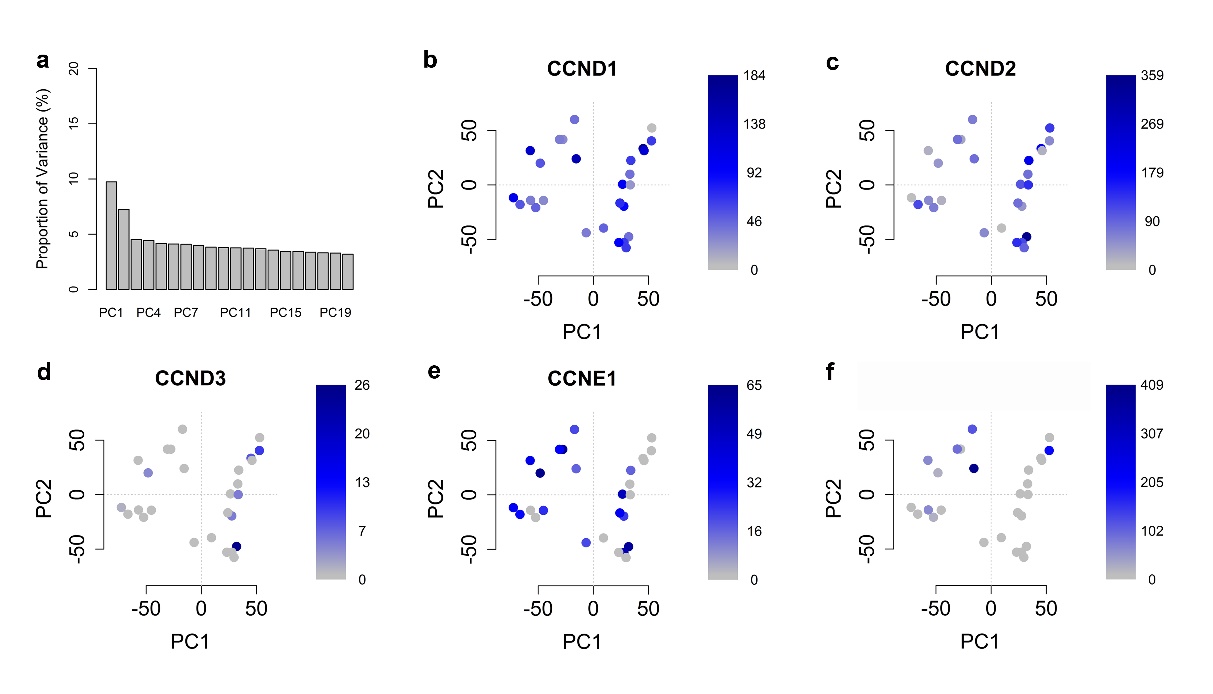

**Extended Data Figure 5 | Genes associated with G1/S phases of the cell cycle appear to be randomly distributed among node cells.** The first two PCs provide the most informative measure for clustering (**a**). The dataset comprises 27 cells collected from the HH8 node, originating from lateral epiblast grafts (as shown in Fig. 3c). The first 20 out of 27 principal components are shown. Expression of a selection of cyclins associated with the G1/S-phases of the cell cycle are represented in (**b-f**). FPKM levels reflected by intensity of blue. For the expression profile of G2/M cell cycle related genes associated with PC1 see ‘Extended Data Figure 4’.

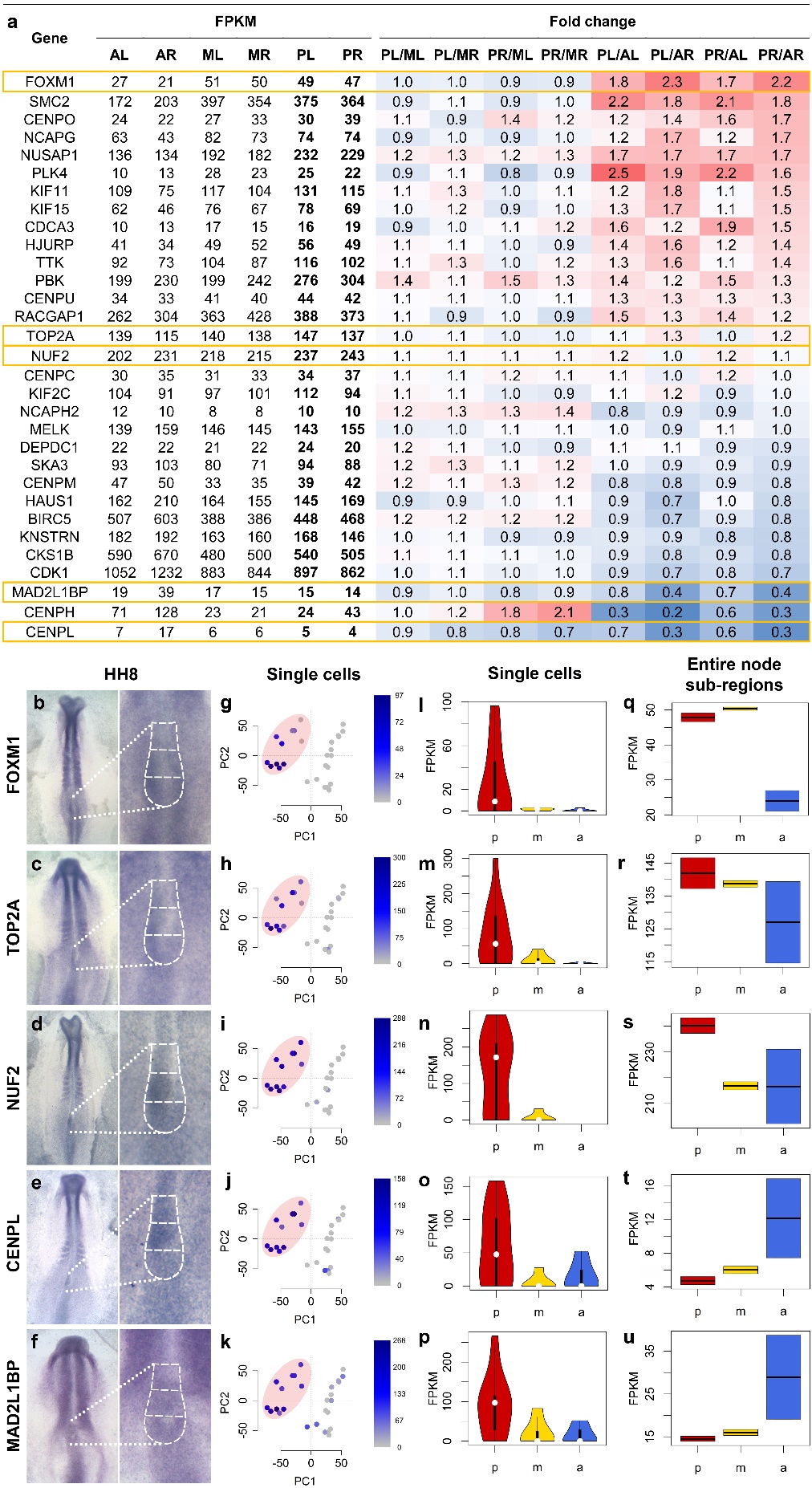

**Extended Data Figure 6 | Single resident cells express G2/M phase cell-cycle related genes in a distinctive manner; for some genes, this is different from the dominant pattern of expression in the same sub-region of the node.** G2/M phase cell cycle related genes were found to be enriched in single cells from the posterior node (Extended Data Figure 4). **a,** In the present table, expression of these G2/M cell cycle related genes is shown in entire node sub-regions obtained by bulk RNA-seq (AL, anterior left; AR, anterior right; ML, middle left; MR, middle right; PL, posterior left; PR, posterior right, and see ‘Extended Data Figure 7a’). Red: fold change >1; blue: fold change <1. Expression of genes highlighted in yellow also assessed spatially by (*in situ* hybridization) (**b-f**), at the single cell level (scRNA-seq, see Fig. 3c) (**g-p**) and then compared to expression in entire node sub-regions (bulk RNA-seq data, see ‘Extended Data Figure 7’) (**q-u**). **b-f,** *in situ* hybridization shown in ventral view. **g-k,** Pink ovals show the cluster containing most cells derived from the posterior part of the node. FPKM levels reflected by intensity of blue. **l-p,** Violin plots show that the highest levels of G2/M phase cell cycle related genes are in single cells from the posterior (p, red) region of the node when compared to the middle (m) and anterior (a) regions. **q-u,** Boxplots show that the expression across entire node sub regions is also higher in the posterior node sub-regions for FOXM1 (**q**), TOP2A (**r**) and NUF2 (**s**), but is instead higher in the anterior node sub-regions for CENPL (**t**) and MAD2L1BP (**u**).

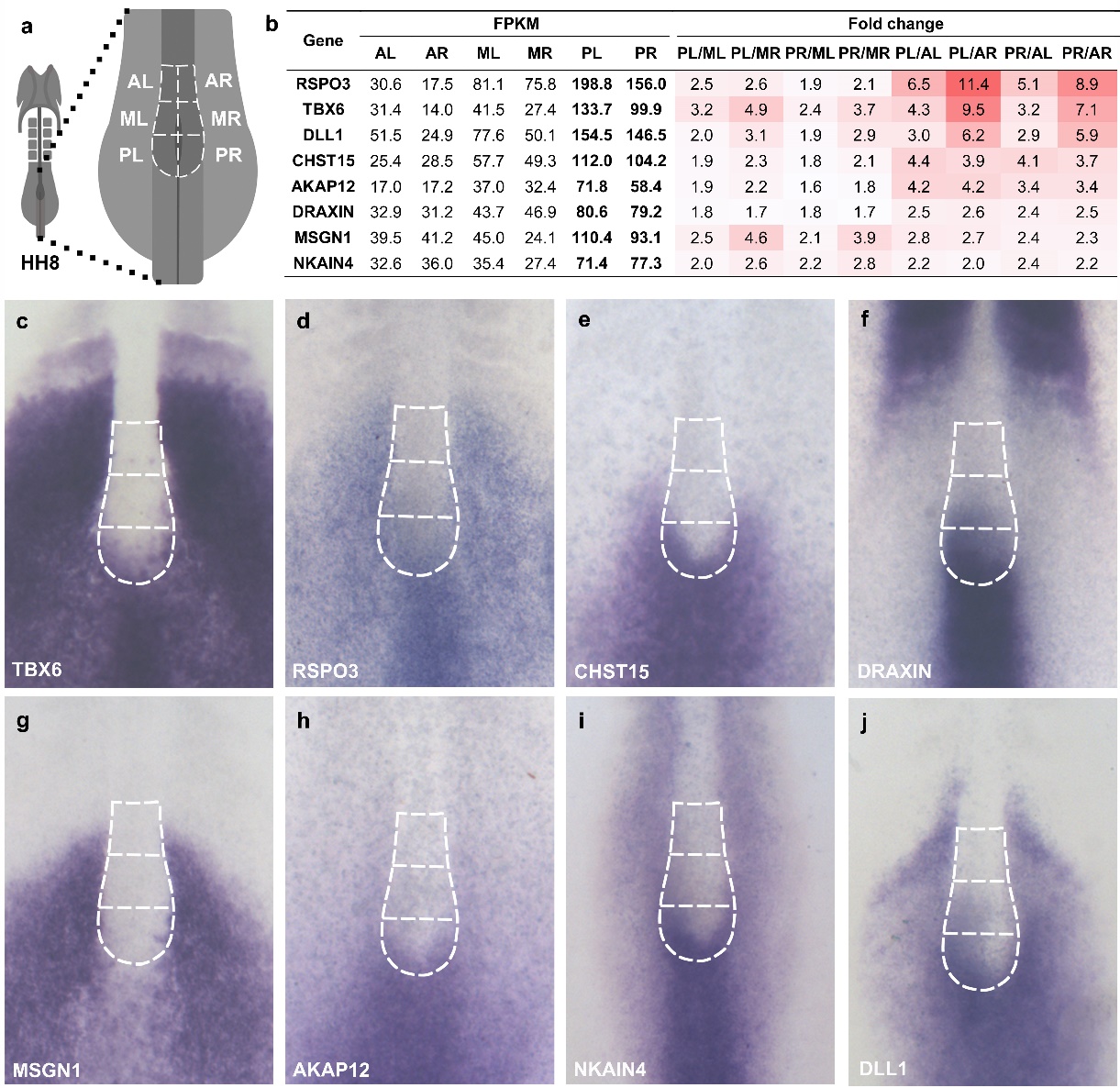

**Extended Data Figure 7 | Examples of genes with higher expression in posterior node, as revealed by RNA-seq of entire node sub-regions, verified by *in situ* hybridization. a,** Node sub-regions sampled for bulk RNA-seq. HH8 node outlined with white-dashed line and sub-divided into six sub-regions (ventral view): anterior left (AL), anterior right (AR), middle left (ML), middle right (MR), posterior left (PL) and posterior right (PR).  **b,** Genes with expression enriched in the posterior node (PL, PR) (data from bulk RNA-seq). All selected genes have a fold-change of at least 1.5 in PL and PR versus AL, AR, ML and MR, and have an FPKM value of at least 50 in PL and PR. The intensity of red corresponds to degree of fold change. **c,** *In situ* hybridization of genes from (**b**). Anterior, middle and posterior regions of the node outlined. All embryos are between HH8- and HH8+, shown in ventral view. The pattern of expression within the posterior sub-region of the node can be ‘salt-and-pepper’ (**c, g**), ‘diffuse’ (**d**), ‘horseshoe-like’ (**e, h, i**) or ‘asymmetrical’ (**f, j**). Several of these genes are characteristic of different signaling pathways (Wnt, FGF and Notch) and/or have been implicated in stem cell niches in other systems (see main text).

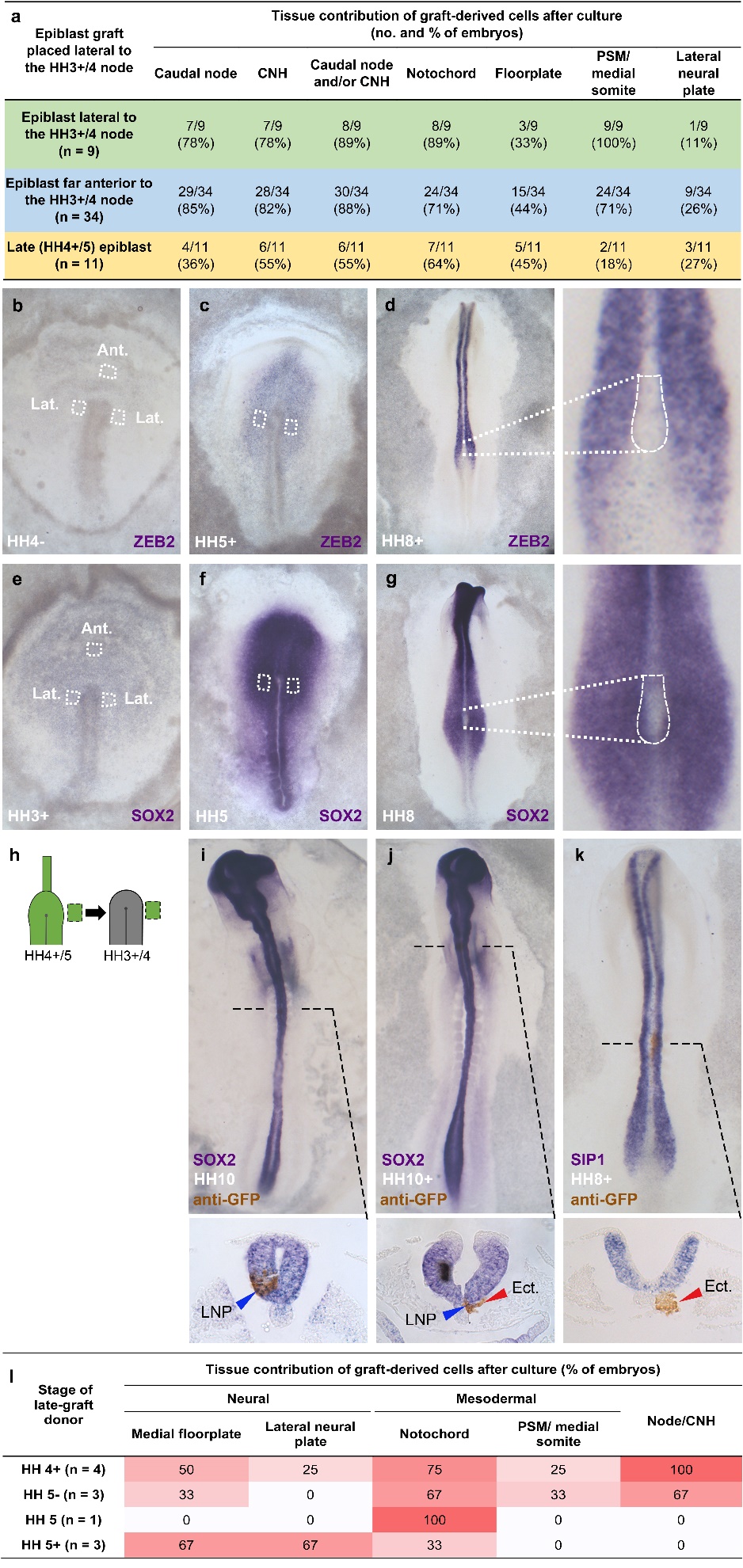

**Extended Data Figure 8 | Late epiblast derived cells only express neural markers when in the neural plate. a,** Table comparing the tissue contributions from epiblast grafts from late (HH4+/5) and early (HH3+/4) donor embryos. HH3+/4 donor epiblast grafts (green and blue) performed as shown in Fig. 1d and f. Late epiblast donor grafts (yellow) performed as in Fig. 4c. n-numbers indicate the number of embryos grafted. **b,** The neural plate markers ZEB2 and SOX2 are highly expressed in late (HH4+/5) donor epiblast but absent or low in early (HH3+/4) donor epiblast. (Ant.: position from which anterior donor epiblast was taken; Lat.: region from which lateral donor epiblast was dissected). Expression in the regressing node is seen only in a few cells (**d, g**). All embryos shown in dorsal view. **h-k,** Late epiblast derived cells only express neural markers when in the neural plate. Late (HH4+/5) epiblast was grafted lateral to the HH3+/4 node (**h**). Resulting embryos after culture, processed for *in situ* hybridization and anti-GFP antibody staining (**i-k**). Whole mounts in dorsal view. The blue arrow indicates GFP-positive cells (brown) that overlap with the *in situ* signal (purple). The red arrow points to GFP-positive cells in ectopic locations not overlapping with the *in situ* signal. **l,** A transition away from mesodermal and towards neural fates in lateral epiblast occurs around stages HH5- to HH5. Epiblast grafts performed as in (**h**). n-numbers indicate the number of grafted embryos. The intensity of red reflects the proportion of embryos with graft-derived cells in the tissues indicated. CNH, chordoneural hinge; PSM, presomitic mesoderm. LNP, lateral neural plate; Ect., cells from the graft that failed to integrate into any host structure.

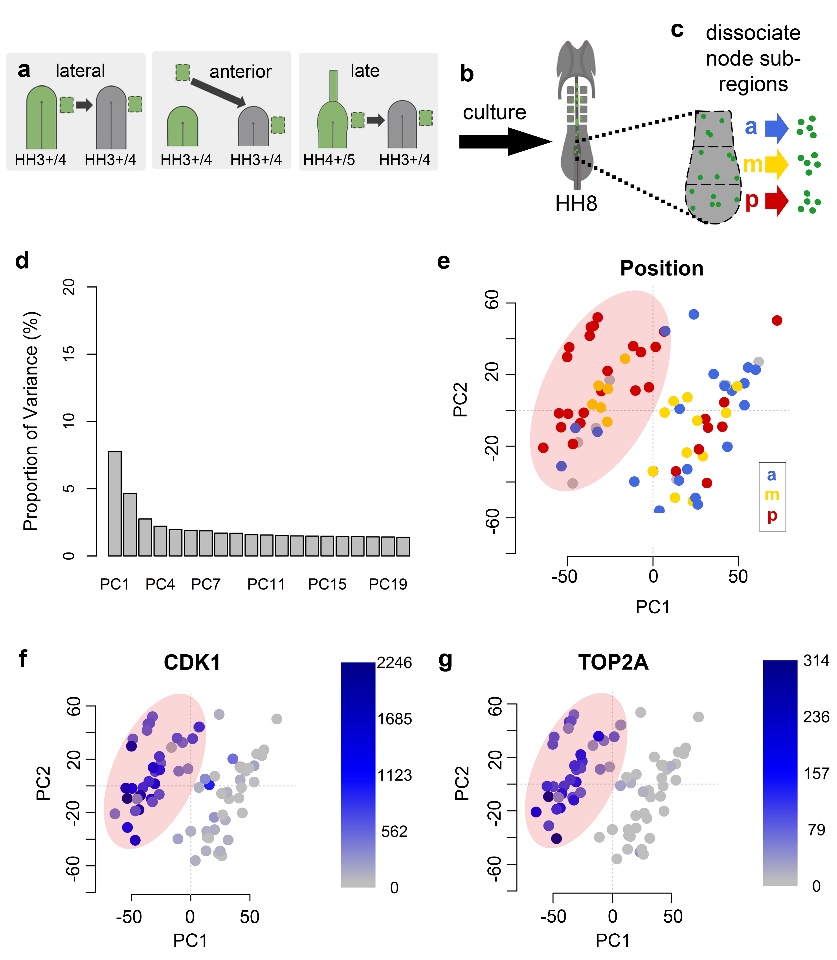
**Extended Data Figure 9 | Irrespective of a cell’s origin, most variation among cells in the node is explained by expression of G2/M phase related cell-cycle genes.** All single cells were collected from the HH8 node but originate from three graft conditions. **a,** Epiblast just lateral to the node that normally enters the node (‘lateral’) and epiblast that was made to enter the node from ‘anterior’ or ‘late’(HH4+/5) epiblast. **b-c,** Grafted embryos were cultured to HH8 (**b**), the node dissociated and single GFP-positive cells isolated from anterior (a, blue), middle (m, yellow) and posterior (p, red) parts of the node and processed for scRNA-seq (**c**). **d,** Principal components (PCs) as a measure of variation in a dataset comprised of 77 cells collected from the HH8 node. The first 20 out of 77 PCs are shown. The first three PCs provide the most informative measure for clustering. **e-g,** Cells plotted according to PC1-2. Pink ovals contain most cells from the posterior part of the node (‘posterior cluster’). The grey spots in (**e**) mark cells whose position could not be ascertained. Expression of G2/M phase cell-cycle related genes limited almost exclusively to cells in the ‘posterior cluster’ (**f-g**) (FPKM levels reflected by intensity of blue).

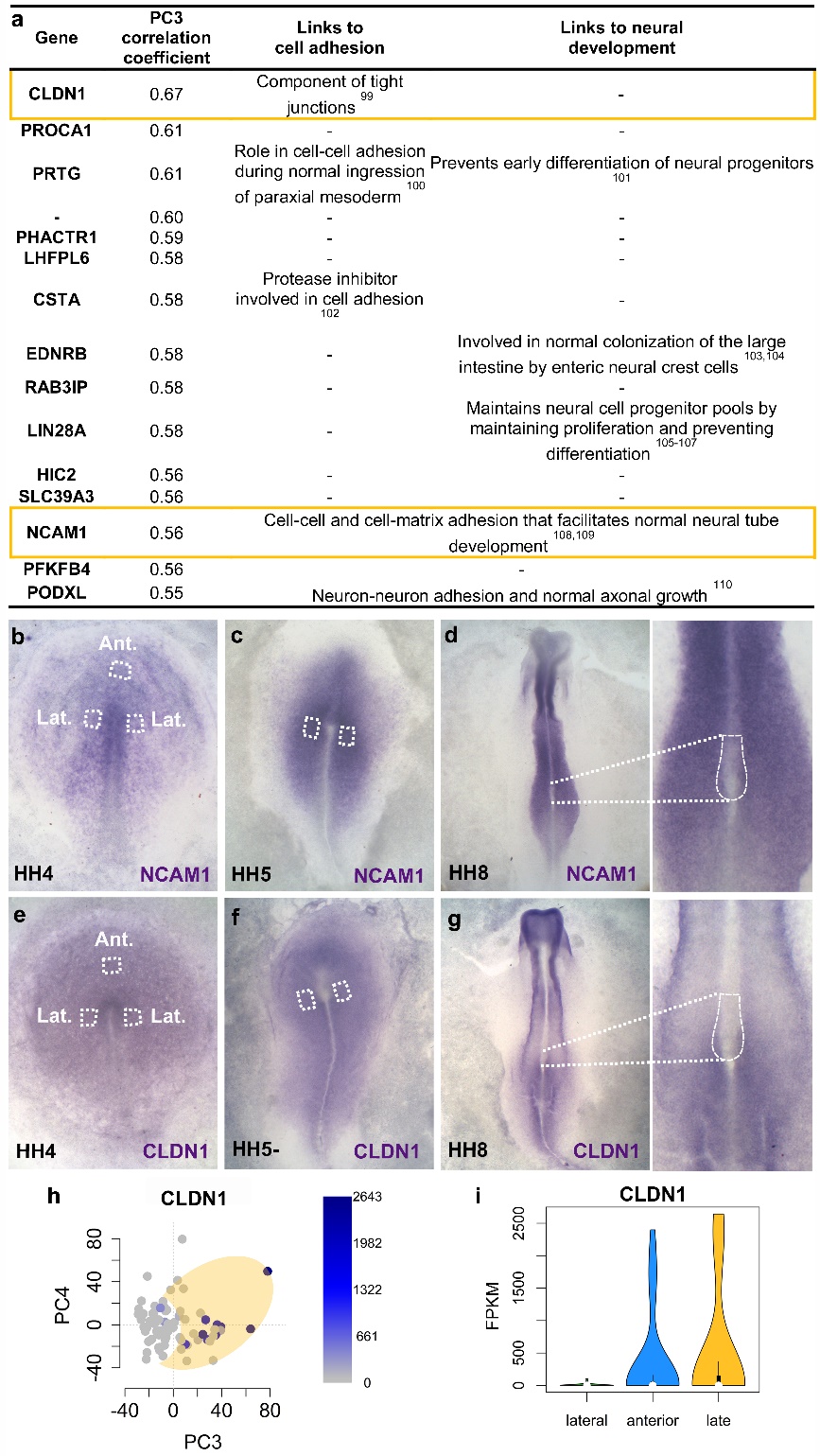
­­

**Extended Data Figure 10 | Cells derived from late epiblast are characterized by a neural-plate-like gene expression signature. a,** Correlation of gene expression with PC3 from scRNA-seq data. Of 15 genes with a correlation coefficient ≥0.55 for PC3, at least 7 are linked to cell adhesion and/or neural development. **b-g,** Expression of genes highlighted in yellow assessed spatially by *in situ* hybridization. NCAM1 (**b-d**) and CLDN1 (**e-g**) are expressed in donor epiblast from early (HH3+/4) (**b, e**) and late (HH4+/5) stage embryos (**c,** **c**) (Ant.: position from which anterior donor epiblast was taken; Lat.: region from which lateral donor epiblast was dissected). These markers are only expressed in a few cells of the node at the time of collection for scRNA-seq (**d, g**). All embryos shown in dorsal view. **h-i,** Expression of CLDN1 among single cells sequenced from the HH8 node correlates with PC3 (**h**) and is restricted to cells originating from late epiblast (yellow) and anterior HH3+/4 epiblast (blue) (**i**) (yellow oval: late epiblast-derived cells not overlapping with the HH3+/4 lateral epiblast derived cells).
